# Microevolution of bank voles (*Myodes glareolus*) at neutral and immune-related genes during multiannual dynamic cycles: Consequences for Puumala hantavirus epidemiology

**DOI:** 10.1101/048082

**Authors:** Dubois Adelaïde, Maxime Galan, Jean-François Cosson, Bertrand Gauffre, Heikki Henttonen, Jukka Niemimaa, Maria Razzauti, Liina Voutilainen, Renaud Vitalis, Emmanuel Guivier, Nathalie Charbonnel

## Abstract

Understanding how host dynamics, including variations of population size and dispersal, may affect the epidemiology of infectious diseases through ecological and evolutionary processes is an active research area. Here we focus on a bank vole (*Myodes glareolus*) metapopulation surveyed in Finland between 2005 and 2009. Bank vole is the reservoir of Puumala hantavirus (PUUV), the agent of nephropathia epidemica (NE, a mild form of hemorrhagic fever with renal symptom) in humans. *M glareolus* populations experience multiannual density fluctuations that may influence the level of genetic diversity maintained in bank voles, PUUV prevalence and NE occurrence. We examine bank vole metapopulation genetics at presumably neutral markers and immune-related genes involved in susceptibility to PUUV (*Tnf*-promoter, *Mhc-Drb, Tlr4*, *Tlr7* and *Mx2* gene) to investigate the links between population dynamics, microevolutionary processes and PUUV epidemiology. We show that genetic drift slightly and transiently affects neutral and adaptive genetic variability within the metapopulation. Gene flow seems to counterbalance its effects during the multiannual density fluctuations. The low abundance phase may therefore be too short to impact genetic variation in the host, and consequently viral genetic diversity. Environmental heterogeneity does not seem to affect vole gene flow, which might explain the absence of spatial structure previously detected in PUUV in this area. Besides, our results suggest the role of vole dispersal on PUUV circulation through sex-specific and density-dependent movements. We find little evidence of selection acting on immune-related genes within this metapopulation. Footprint of positive selection is detected at *Tlr-4* gene in 2008 only. We observe marginally significant associations between *Mhc-Drb* haplotypes and PUUV serology, and between *Mx2* genotype and PUUV genogroups. These results show that microevolutionary changes and PUUV epidemiology in this metapopulation are mainly driven by neutral processes, although the relative effects of neutral and adaptive forces could vary temporally with density fluctuations.

## 1. Introduction

Molecular epidemiology has largely focused on the parasite side, contributing to a better understanding of the sources of infection, routes of transmission and factors influencing disease diffusion or pathogenesis. This approach remains less applied to hosts, although it can bring insights into factors influencing pathogen persistence, spread and evolution (Archie et al., 2009). In particular, host population genetics may enable to infer major demographic characteristics, including population size and individual movements, which play critical roles in epidemiology. Besides, many parasites lack free-living stages or have limited dispersal capacities and are therefore highly dependent of their hosts to disperse (Price, 1980). When disease transmission is specifically associated with a given host species, host population genetics can reveal environmental heterogeneity that may affect the spread of diseases by reducing both movements of infected hosts and contacts between infected and non-infected ones (Biek and Real, 2010). Landscape genetics studies in the host could therefore help predicting this spread of diseases and developing proactive disease management. Analysing the adaptive genetic variation that is linked to host–pathogen interactions can also improve our knowledge of disease epidemiology (see for a review Charbonnel and Cosson, 2011). Molecular epidemiology has enabled to identify host genes that are related with host susceptibility and resistance to infectious diseases. This information is essential to better assess the number of susceptible individuals within host populations. Although these genes are expected to strongly evolve under directional pathogen-driven selection, high levels of polymorphism have been observed in natural populations. This is mainly due to the interplay of neutral and selective microevolutionary forces (Charbonnel and Cosson, 2011). Because populations are ususally not at equilibrium and experience demographic events such as bottlenecks, a decline in the efficiency of selection might for example be observed at these immune-related genes (Ejsmond and Radwan, 2011; Grueber et al., 2013; Sutton et al., 2011). Besides, in particular cases including genes of the Major Histocompatibility Complex (*Mhc*), balancing selection, in addition to migration, may rapidly buffer the loss of adaptive diversity in bottlenecked populations (Oliver and Piertney, 2012; Winternitz et al., 2014).

Multiannual population density cycles are extreme cases of non-equilibrium demographic situations that could benefit from the development of host population genetics to address epidemiological questions. Such dynamics can lead to variations of effective population size (Frankham, 1995) that may significantly affect genetic diversity and population genetic structure. Drastic reductions in population size can cause a rapid decrease of allelic richness and a gradual erosion of heterozygosity, with negative consequences on adaptive potential (Bryja et al., 2007; Maruyama and Fuerst, 1985; Nei et al., 1975). However a high level of genetic diversity may sometimes persist both at neutral and immune-related genes (Berthier et al., 2006; Burton et al., 2002; Ehrich et al., 2009; Gauffre et al., 2014; Rikalainen et al., 2012; Winternitz et al., 2014). This phenomen is mainly explained by a metapopulation functioning with small demes experiencing extinction-recolonisation during low density phase and spatial expansion with an increase of the effective migration during high abundance phase. As such, the genetic diversity is shown to remain relatively constant through time when considering all demes together (Berthier et al., 2006; Burton et al., 2002; Gauffre et al., 2014; Rikalainen et al., 2012; Winternitz et al., 2014). Density cycles can also impact dispersal rates, and consequently the distribution of genetic variability within and among populations. Several studies showed that cyclic populations of the fossorial water vole were weakly differentiated during the increasing phase of abundance but highly differentiated during the low phase of abundance, as a result of local genetic drift and low dispersal rates (Berthier et al., 2006, 2005). Such cyclic dynamics could therefore have severe consequences on disease epidemiology, both ecologically and evolutionarily. Up to now, most of the studies dedicated to the genetics of these cyclic populations have been done to infer demographic characteristics and to analyse changes in immune-related genes polymorphism. The links with epidemiological risks and host-pathogen interactions still remain to be explored.

In this study, we examined this tripartite relationship between host density cycles, population genetics and epidemiology of a viral infection in the bank vole (*Myodes glareolus*) and Puumala hantavirus (PUUV, family of *Bunyaviridae*) as model species. In boreal Fennoscandia, bank vole populations undergo three to five year density cycles with high amplitude (Hanski and Henttonen, 2002; Henttonen et al., 1985). The bank vole is the reservoir of PUUV, the etiologic agent of nephropathia epidemica (NE) (Vaheri et al., 2012). Among voles, the virus is transmitted horizontally through direct contacts between individuals (*i.e*. bites) or indirectly via infectious aerosolized urine and feces (Bernshtein et al., 1999; Kallio et al., 2006a). The spread of PUUV is therefore closely linked to its host dispersal. PUUV infection in bank voles is chronic and mainly asymptomatic (Bernshtein et al., 1999; Meyer and Schmaljohn, 2000; Voutilainen et al., 2015) as maturation, winter survival and female breeding of infected individuals may be affected (Kallio et al., 2015, 2007; Tersago et al., 2012). The epidemiology of PUUV in rodent populations has been well documented (Voutilainen et al., 2016). Only two studies have yet examined the connection between bank vole population and genetic microevolution of PUUV. Guivier et al. (2011) developed a landscape genetics approach in temperate zone to analyze the influence of bank vole population dynamics on PUUV spatial distribution in contrasted landscape structures. They considered genetic diversity as a proxy for estimating vole population size, isolation and migration, and found that it was highly positively correlated with PUUV prevalence and with the abundance of forest habitats at a local scale. This suggested that the persistence of PUUV was enhanced in lowly fragmented forests, where vole metapopulations are highly connected and experience low extinction rates. This result was congruent with previous correlations found between human PUUV incidence and coverage of beech tree forests (Clement et al., 2010). Besides, Weber de Melo et al. (2015) made a comparative analysis of the spatiotemporal population genetic structure of bank voles and PUUV at a smaller spatial scale. They aimed at testing the influence of vole neutral microevolutionary processes on PUUV phylodynamics. They found incongruent patterns between host and PUUV population genetic structure that were likely to arise due to the high evolutionary rate of this virus.

What is still missing to better understand the links between bank vole metapopulation dynamics and PUUV epidemiology is the characterization of the microevolutionary processes at play in the genes known to influence rodent susceptibility to hantaviruses (review in Charbonnel et al., 2014). To bridge this gap, we analyzed the genetics of bank voles at presumably neutral and PUUV associated genes (namely *Mhc-Drb, Tlr4* and *Tlr7, Tnf-α* promoter and *Mx2* immune-related genes) in a metapopulation monitored during a whole demographic cycle in an highly endemic NE area of Finland. We first examined the consequences of drastic density variations on the dynamics of neutral and immune-related genetic diversity in *M. glareolus* populations, both at the metapopulation scale and at the ‘deme’ scale. Second, we performed population genetics analyses to look for spatial barriers, like lake or rivers that could affect gene flow within the bank vole metapopulation and therefore the circulation of PUUV. We also tested whether PUUV transmission was likely to result from vole dispersal or contacts between relatives (Deter et al., 2008). Third, we investigated the possibility that selective forces may shape bank vole / PUUV interactions despite the pronounced multiannual density cycles. To this aim, we compared the genetic structure observed at immune-related genes to that measured at presumably neutral microsatellite markers, assuming that differences between the two could be the consequence of selection. We furthermore looked for associations between immune-related gene polymorphism, susceptibility to PUUV and PUUV genetic variants.

## 2. Materials and Methods

### 2.1. Ethic statement

All handling procedures of wild live bank voles followed the Finnish Act on the Use of Animals for Experimental Purposes (62/2006) and took place with permission from the Finnish Animal Experiment Board (license numbers HY 45-02, HY 122-03, and HY 54-05). All efforts were made to minimize animal suffering. The species captured for this study, *Myodes glareolus*, neither is protected nor included in the Red List of Finnish Species. The animal trapping took place with permission from the landowners.

### 2.2. Rodent sampling and PUUV detection

Rodents were trapped within an area of about 25 km² of typical spruce-dominated taiga forest at Konnevesi, Central Finland (62° 34’ N, 26° 24’ E), where PUUV is highly endemic (Kallio et al., 2007; Razzauti et al., 2013, 2008; Voutilainen et al., 2015; Voutilainen et al., 2016). Trappings for bank vole genetic materials were carried out between 2005 and 2009 to include a whole density cycle (peak year 2005, crash year 2006, increase 2007, peak 2008, decline 2009, see (Razzauti et al., 2013; Rikalainen et al., 2012; Voutilainen et al., 2015), Fig. 1). Too few individuals could be sampled in crash year 2006 so that individuals sampled in 2006 were not included in the study. Trapping sessions were conducted three times per year, in the beginning and middle period of the breeding season (May, July in 2005, 2008, 2009) and in October (2007, 2009) to gather enough individuals when abundance levels were low. Nine sites situated at least 750m apart from each other within 25 km^2^ were sampled (Fig. 1). Briefly, a grid of 3 × 3 Ugglan Special live traps (Grahnab AB, Sweden) with 15 m intervals and baited with oat seeds and potato was set in each site. Traps were checked once a day during three days, and captured bank voles were brought into the laboratory, bled through retro-orbital sinus, sacrificed, weighed, measured and sexed (for details, see Voutilainen et al. (2016)). Tissue samples (toe or tip of the tail) were collected and stored in 95% ethanol for further genetic analyses. The heart of each animal was collected, diluted into 200 µL phosphate-buffered saline and frozen for latter PUUV antibody analysis. Detection of PUUV was done as described in Kallio-Kokko et al. (Kallio-Kokko et al., 2006). Serological and virological data have previously been described in Voutilainen et al. (2016, 2015) and Razzauti et al. (2013, 2008). Because PUUV infection in bank voles is chronic, the presence of PUUV antibodies was taken as an indicator of infection (Meyer and Schmaljohn, 2000). Yet, because juvenile voles trapped in July could be false positive because of the presence of maternal antibodies (Kallio et al., 2006b), all bank voles with low body mass (<17.5g, see Voutilainen et al. 2016) were considered juveniles carrying maternal antibodies but not PUUV infected (21 bank voles were concerned and were not considered in the analyses including seroprevalence). Sampling size and seroprevalence levels are described in details in Table S1.

**Fig 1.**
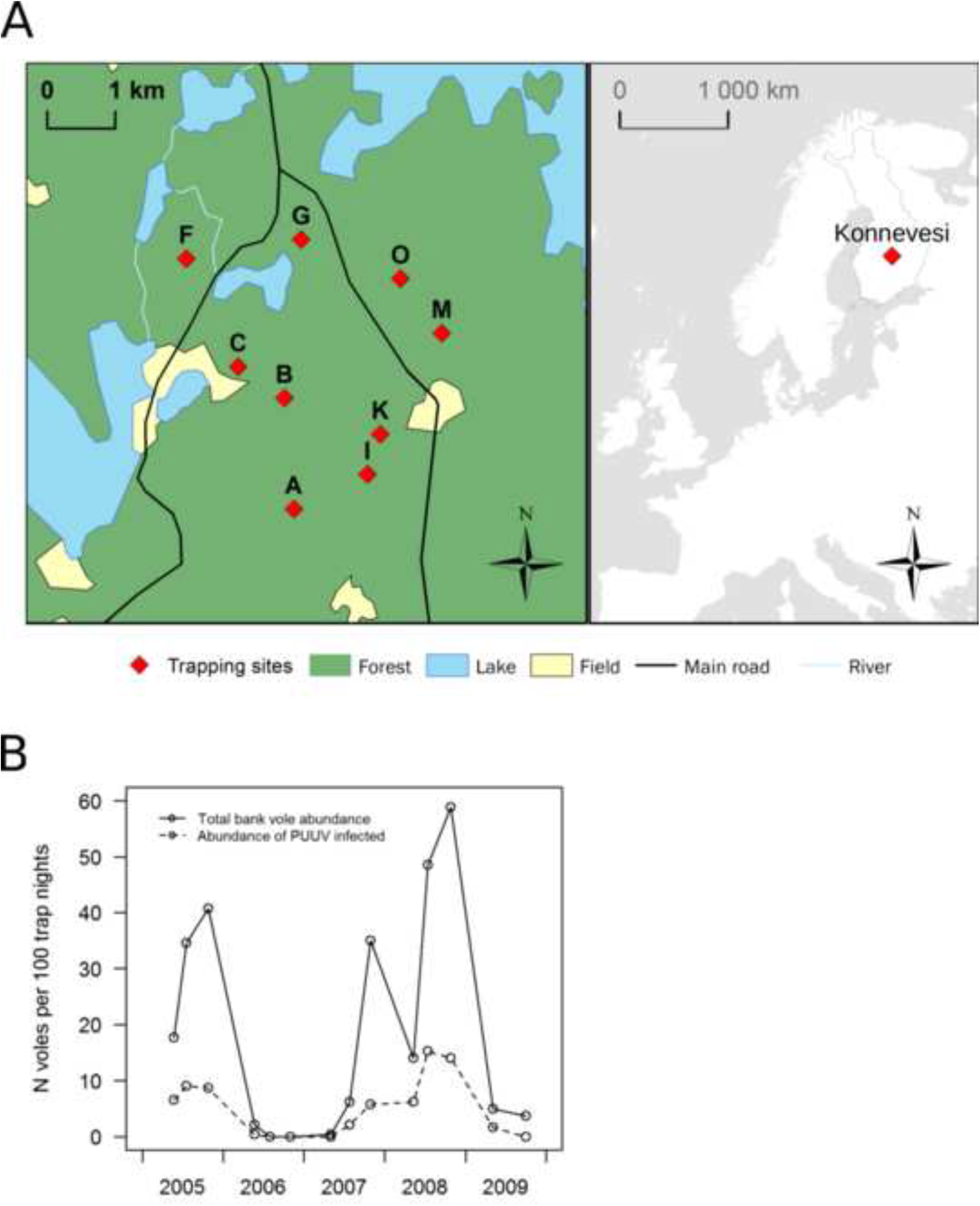
Map of the study area and long-term fluctuations of bank voles in Konnevesi. A) The nine trapping sites included in this study are indicated by a red square. These sites were situated at least 750 m apart. The location of Konnevesi within Finland is shown on the right box. B) Total bank vole abundance and number of infected voles trapped per 100 trap nights in the nine Konnevesi sites considered in this study between 2005 and 2009 (from (Voutilainen et al., 2016)). White circles represent sampling dates. This figure contains data from the National Land Survey of Finland Topographic Database 06/2013 (left panel) and Esri, HERE, and DeLorme (right panel).

### 2.3. DNA extraction and loci amplification

*Microsatellite analyses* –Genomic DNA was extracted using the DNeasy 96 Tissue kit (Qiagen) according to manufacturer’s recommendations. A final elution was done in 400μl of AE buffer (Qiagen). Samples were first genotyped at 19 unliked microsatellites loci previously published by Rikalainen et al. (Rikalainen et al., 2008) using primers and cycling conditions described in Guivier et al. (Guivier et al., 2011). Genotyping was carried out using an ABI3130 automated DNA sequencer (Applied Biosystems). Alleles were scored using GENEMAPPER software (Applied Biosystems). Two loci were excluded from the genetic analyses: Cg6G11 because of the poor quality profiles obtained and Cg5F11 because it was duplicated in these populations.

*Candidate gene analyses* - The polymorphism of candidate genes that are relevant with regard to PUUV infections was further analyzed. These candidate genes, namely *Mhc-Drb, Tlr4* and *Tlr7, Tnf-α* promoter and *Mx2* were selected based on the literature of rodent or human/hantavirus interactions (Charbonnel et al., 2014; Rohfritsch et al., 2013). The second exon of the *Mhc-Drb* class II is a highly polymorphic gene of the major histocompatibility complex. It encodes for proteins responsible for pathogen recognition (Klein and O’Huigin, 1994). *Tlr4* and *Tlr7* genes encode for Toll-like receptors which play an important role in activating the innate immune system (Ayres and Schneider, 2012). TLR4 is a universal receptor able to detect different ligands as LPS of Gram-negative bacteria and also some viral proteins (Wlasiuk and Nachman, 2010) including hantaviral proteins (Sabeti et al., 2007). TLR7 is a receptor to virus, which is probably involved in the detection of ssRNA viruses like hantaviruses (Bowie and Haga, 2005). As both *Mhc* and *Tlr* genes encode for receptors, their polymorphism is likely to be linked to the efficiency of antigen recognition. *Tnf* (Tumour Necrosis Factor alpha) gene encodes for a cytokine involved in the pro-inflammatory response (Vassalli, 1992). Polymorphism within the *Tnf* promoter seems to be, at least partly, implicated in the regulation of *Tnf* transcription and consequently in the magnitude of the secretory response of this cytokine (Bouma et al., 1996; Wilson et al., 1997). Mx proteins are interferon-induced members of the superfamily of large GTPases. They mediate innate, antiviral resistance in rodents, as well as in other vertebrates and humans (Jin et al., 1999; Porter et al., 2006). Most Mx GTPases inhibit a wide range of RNA viruses, including members of the *Bunyaviridae* at an early stage in their life cycle. Thus, the efficiency of antiviral resistance seems to be mediated by *Mx* genes polymorphism.

The second exon of the *Mhc-Drb* class II was genotyped using a 454 Genome Sequencer FLX (Roche) with the procedure detailed in Galan et al. (2010). The locus was amplified using a modified version of the primers JS1 (5’- GCCTCCCTCGCGCCATCAG −3’) and JS2 (5’- GCCTTGCCAGCCCGCTCAG −3’) initially designed for a lemur (Schad et al., 2004). Individual-specific MIDs (Multiplex Identifiers) and 454 adaptors required for the emPCR and the 454 GS-FLX sequencing were also added (for more details concerning laboratory procedures, see Galan et al., 2012, 2010). The SESAME barcode software (SEquence Sorter & AMplicon Explorer (Piry et al., 2012)) was used to sort sequences, identify and discard artefactual variants, and generate the allele sequences and individual genotypes. The *Drb*-exon2 sequences were aligned in the BioEdit sequence alignment editor (Hall, 1999) using CLUSTAL X Multiple Alignment v7.0.9.0 (Chenna et al., 2003). Nomenclature of the bank vole sequences follows Klein et al. (1993). The sequences are available in GenBank under accession numbers HM347503–HM347507 and HM107849– HM107870. Human *Homo sapiens* (GenBank accession numbers 91M17236 and AM109973), sheep *Ovis aries* (M33304, AY230000) and pig *Sus scrofa* (M29938, NM_001113695) sequences were used as outgroups in phylogenetic analyses. MEGA4 software (Tamura et al., 2007) was employed to construct a phylogenetic tree of the *Drb*-exon2 gene using the neighbor-joining algorithm and Kimura two-parameter distance model. A bootstrap analysis (1000 replicates) was performed to determine the reliability of the branching. As previously described in Axtner & Sommer (2007) and observed in Guivier et al. (2010) and Rohfritsch et al. (2013), one to six different haplotypes could be observed for a single bank vole, indicating the presence of duplications in the *Mhc-Drb*-exon2 gene, the number of which varies between individuals. The phylogenetic reconstructions did not enable us to assign all haplotypes to a particular copy of the *Mhc-Drb* gene (Supplementary Material Fig. S2). However, we confirmed that cluster I, first described by Axtner & Sommer (2007), is highly supported (see also Kloch et al., 2010; Scherman et al., 2014). We could therefore identify unambiguously the different haplotypes belonging to this monophyletic cluster. We also verified that each bank vole carried two haplotypes from cluster I at most, what highly corroborated the fact that it corresponds to a single *Mhc-Drb* gene copy. As it was not possible to delineate any other cluster, the remaining haplotypes were not considered in further population genetics analyses.

Polymorphisms of *Tnf* gene promoter (Guivier et al., 2010), *Tlr4* and *Tlr7* gene, and *Mx2* exons 13 and 14, were previously investigated by the sequencing of bank voles sample from several European countries. Protocol S1 and Table S2 provide details on the list of primers and PCR conditions used to amplify the gene products. At the European scale, we identified ten single nucleotide polymorphisms (SNPs) in *Tlr4* exon 3, one SNP in *Tlr7* exon 3 and four SNPs in *Mx2* exons 13 and 14 (Table S2, Charbonnel et al., 2014; Rohfritsch et al., 2013). Based on this previous pilot study, we selected the eight polymorphic SNPs that were not singletons (haplotype frequencies higher than 5%, (Bollmer et al., 2011; Quéméré et al., 2015). Two of them were non-synonymous. The software PolyPhen (Ramensky et al., 2002) was used to predict the functional impact of these non-synonymous SNPs on the translated protein, based on biochemical and physical characteristics of the amino acid replacements. The 382 individuals from Konnevesi were genotyped at these SNPs using the robust KASPar allelic-specific fluorescent genotyping system provided by KBiosciences (Hoddesdon, UK).

### 2.4. Statistical analyses

*Within site and within metapopulation genetic diversity* – As the number of voles trapped per site could be low, we pooled samples from May and July for each site. Moreover, because 2009 was a low-density year, we pooled all samples from all sites between May and October for this year. Further analyses were realized at two different spatial scales for each year: for each site and over the whole metapopulation. Considering microsatellites, we tested the conformity to Hardy-Weinberg equilibrium (HWE) for each locus and we analyzed linkage disequilibrium (LD) for each pair of loci using GENEPOP v4.2 (Raymond and Rousset, 1995). Corrections for multiple tests were performed using the false discovery rate (FDR) approach as described in Benjamini & Hochberg (1995). For all microsatellites and for each immune gene (*Mhc-Drb, Tnf, Mx2*, *Tlr4* and *Tlr7*), we estimated *N*_a_, the estimate of allelic richness corrected for minimum sample size (8 in our case) (El Mousadik and Petit, 1996; Goudet, 1995), the observed (*H*_o_) and Nei’s unbiased expected (*H*_e_) heterozygosities (Nei, 1978) and Weir & Cockerham’s inbreeding coefficient *F*_IS_ (Weir and Cockerham, 1984). Significance of *F*_IS_ (excess or deficit in heterozygotes) was assessed using 1000 permutations of alleles within each site. Similar estimates were obtained over the whole metapopulation. Calculations and tests were realized using GENETIX v4.05.2 (Belkhir et al., 1996), using FDR corrections. For these analyses, the (*year*site*) combinations with less than eight individuals were not considered in order to avoid statistical biases.

To test whether these indices of within site and metapopulation genetic diversity varied throughout population dynamic cycles, we defined two linear mixed models, using the LME() function implemented in the NLME package for R 3.1.0 (R Core Team, 2013). Model validations were realized *a posteriori* by checking plots of the residuals. The first model included independently the indices of diversity *N*_a_ and *H*_e_ estimated for the sites sampled in 2005, 2007 and 2008 (2009 was not included as sample sizes per site were too low). Fixed effects included the variables *year*, *site*, *type of loci* (microsatellites, SNPs, *Mhc-Drb*). We also included the pairwise interactions (*year*type of loci)* and (*year*site)*. When the pairwise interactions were non-significant, they were removed from the model. The variable *loci* was included as a random effect to reflect the fact that there were initial differences in diversity between loci. The second model included the global *N*_a_ and *H*_e_ indices estimated over the whole metapopulation, for each year between 2005 and 2009 as response variables. The fixed effects *year*, *type of loci,* their pairwise interaction (*year*type of loci)* and the random effect *loci* were also added. For both models, significance of the fixed effects was assessed with analysis-of-deviance tables (function ANOVA). Chi-square tests with Bonferroni correction were then applied to analyze the effect of significant variables using the package phia for R.

*Test for selective neutrality at microsatellite loci –* We used the software package FDIST2 (Beaumont and Nichols, 1996) to test for homogeneity in differentiation level across the final set of microsatellite loci (15 markers, see the Results section). This method is based on the principle that genetic differentiation among populations (as measured by *F*_ST_) is expected to be higher for loci under divergent selection, and lower for loci under balancing selection, as compared to the rest of the genome. The rationale of FDIST2 is to compute *F*_ST_ and the overall heterozygosity at each marker (computed as the average pairwise difference between all possible pairs of genes between each sample), and then to compare these values to a null distribution, generated by means of neutral coalescent simulations in a symmetrical island model at migration-drift equilibrium. Since the method was applied on microsatellite data, we used a stepwise mutation model, as implemented in FDIST2. One million simulations were performed for each analysis, assuming a 25-demes island model (and the same number of sampled demes as in the original dataset). Simulations were performed for each year independently. As in Ségurel et al. (2010), we used the averaged shifted histogram (ASH) algorithm (Scott, 2002) to characterize the probability distribution, to provide a “high probability region” that contains 90%, 95%, or 99% of the total probability distribution, and to derive an empirical *p*-value for the joint (*F*_ST_, *H*_e_) estimates at the candidate genes.

*Spatial genetic structure –* We analysed the genetic differentiation between each pair of sites using microsatellite data and Weir & Cockerham’s pairwise *F*_ST_ estimates (Weir and Cockerham, 1984). Temporal differentiation was evaluated by estimating *F*_ST_ values between sampling dates. We used exact tests implemented in GENEPOP v4.2 (Raymond and Rousset, 1995) to test for differentiation. FDR corrections were applied as described above. We next used analyses of molecular variance (AMOVA) implemented in ARLEQUIN v3.5.1.2 (Excoffier and Lischer, 2010) to quantity the proportions of genetic variation due to differences between years and between sites, using two hierarchical models (see Table 1 for details).

Spatial genetic clustering was performed using microsatellites markers in a single analysis including all years and sites to detect potential environmental factors affecting gene flow. We used the Bayesian clustering method implemented in STRUCTURE (Pritchard et al., 2000), using the admixture model with correlated allele frequencies. We performed 20 independent runs, varying the number of clusters from *K* = 2 to *K* = 8, using a Markov chain Monte Carlo (MCMC) made of 10^6^ iterations following a burn-in of 500,000 iterations. To determine the number of clusters (*K*) best fitting our multilocus genetic data set, we used the method described in Evanno et al. (2005). We next used the FullSearch algorithm implemented in CLUMPP v1.1 (Jakobsson and Rosenberg, 2007) to assign individuals to clusters. Then, summary bar plots were built using DISTRUCT v1.1 (Rosenberg, 2004).

We finally examined the fine-scale spatial pattern of microsatellite genetic variation using inter-individual relatedness. We estimated the *Li* coefficient (Li et al., 1993) implemented in SPAGeDI v1.4.c (Hardy and Vekemans, 2002) for each year, considering each site separately. This estimator shows a high accuracy (low bias) combined with a high precision (low variance) compared to others (Vekemans and Hardy, 2004). We performed spatial autocorrelation analyses to examine how genetic similarity between pairs of individuals changed with geographical distance. This approach was applied independently for each year, and separately for males and females to detect potential differences in dispersal behavior along cycle phases (Gauffre et al., 2014; Rikalainen et al., 2012).

*Links between genetic structure and PUUV epidemiology* - We first tested whether relatedness differed among seropositive and seronegative bank voles, by permuting individuals between these two classes. The seropositive juveniles presumably carrying maternal antibodies were removed from this analysis. Because pairwise data are not independent, we could not directly compare relatedness estimates between these classes of individuals using standard statistical tests (Prugnolle and Meeûs, 2002). We thus applied the procedure of Coulon et al. (2006) that consists in generating 1,000 random resampling sets without replacement, such that each individual occurs only once in a given resampled set. The difference of mean relatedness *D*_*i*_ between the two categories tested is then calculated for each resampling set. Under the null hypothesis, *D*_*i*_ should follow a normal distribution centered on 0. Besides, on one hand, PUUV seropositive bank voles could be less related than seronegative ones if PUUV transmission mainly occurs during dispersal. On the other hand, the opposite pattern is expected if PUUV transmission occurs during winter family clustering, but up to now, this pattern has only been observed in temperate zones (see Hansson, 1986). Therefore we made no *a priori* assumption on the sign of the difference *D*_*i*_ and thus computed two-sided tests.

Second, we evaluated the level of genetic isolation of each site relatively to the others using microsatellite markers and further analysed the links between local genetic characteristics and PUUV seroprevalence. Site-specific *F*_ST_ values and their 95% confidence interval were estimated using the software GESTE (Foll and Gaggiotti, 2006). We then performed a logistic regression with the GLM() function implemented in R. PUUV seroprevalence was the response variable. We defined the explanatory variables as the two first principal components of a principal component analysis (PCA) of the genetic parameters (*H*_*e*_, *H*_*o*_, *F*_*ST local*_, *F*_*IS*_, *N*_*a*_) estimated for each site and each year.

*F*_ST_ *outlier test to detect signatures of selection in candidate genes* – We evaluated whether the genetic differentiation at immune-related genes (*Mhc-Drb* and SNPs) departed from the null distribution expected at neutrality. The rationale was to compute *F*_ST_ and the overall heterozygosity at each immune-related gene, and then to compare these values to a null distribution, obtained by means of neutral coalescent simulations in a symmetrical island model at migration-drift equilibrium. These simulations were generated conditionally on the overall *F*_ST_ estimate calculated from the microsatellite markers only (*F*_ST_ = 0.0209 in 2005; *F*_ST_ = 0.0182 in 2007 and *F*_ST_ = 0.0252 in 2008). Therefore, the null distribution is independent from the level of differentiation observed at the candidate genes using an infinite allele model. For the analysis of the *Mhc-Drb* data, we considered each haplotype as an allele, and we used FDIST2 (Which is specifically designed for the analysis of multiallelic data) to test for departure from the neutral distribution. For the analysis of the SNP data, we used a version of the software package DFDIST, which we modified to generate bi-allelic co-dominant data (Ségurel et al., 2010). An infinite allele model of mutation was considered, with *θ* = 2*nNμ* = 0.1 (where *n* = 25 is the number of demes of size *N*, and *μ* is the mutation rate) and only the simulations resulting in bi-allelic data were retained. One million simulations were performed for each analysis, assuming a 25-demes island model (and the same number of sampled demes as in the original dataset). For all simulations, the maximum frequency of the most common allele allowed was set to 0.99. These analyses were realized for each year independently. We characterized the “high probability regions” (based on the the level of differentiation at neutral markers) and the empirical *p*-values at the candidate genes using the averaged shifted histogram (ASH) algorithm (Scott, 2002), as in Ségurel et al. (2010). Because the true demographic history of *M. glareolus* populations is likely to depart from the underlying assumptions of FDIST2 and the modified DFDIST, especially with respect to the hypothesis of stable demography, we investigated to what extent the approach to detect signatures of selection was robust to temporal variations in population size (see Supplementary Appendix S1).

*Association between genotypes at immune-related genes and PUUV serology* - Because candidate gene polymorphism was expected to be related to bank vole susceptibility to PUUV, we searched for associations between immune variations and PUUV serology which was used as a proxy of PUUV infection status (Guivier et al., 2010). This analysis was realized on the whole dataset, from which we excluded the seropositive juveniles presumably carrying maternal antibodies. First, we performed a correspondence analysis (CoA) on the genetic matrix of candidate gene polymorphism (*Mhc-Drb*: haplotypes; other candidate genes: SNPs, coded as presence/absence), using the software ADE-4 (Thioulouse et al., 1997). Then, we realized a discriminant function analysis (DA) using PUUV serology as a discriminant factor. The significance of the analysis was estimated by using 10,000 permutation tests. The relative risk (RR) associated with each allele of interest, as detected with DA, was estimated following (Haldane, 1956). The significance of the association between these alleles and vole serological status were analyzed using Fisher exact tests and Bonferroni sequential corrections. Similar analyses were performed for seropositive bank voles (*N* = 46) to look for potential associations between PUUV diversity (PUUV genogroups, see Razzauti et al. 2013, 2008) and bank vole immune-related gene polymorphism.

## 3. Results

### 3.1. Validation of microsatellite marker and candidate immune-related gene polymorphism

Two microsatellite loci, Cg2F2 and Cg13F9, showed important deviations from HWE suggesting the presence of null alleles. They were removed from the analyses. The analyses performed using FDIST2 revealed that none of these microsatellites departed from neutral expectation (Supplementary Material Fig. S1). Thus, our final dataset included 382 bank voles genotyped at 15 microsatellites loci.

The 454 sequencing of *Mhc-Drb* exon 2 revealed 17 haplotypes belonging to the monophyletic allele cluster I (Axtner and Sommer, 2007), (Supplementary Material Fig. S2). We did not observe more than two haplotypes from this cluster per individual, what corroborates the fact that these genotypes corresponded to a unique *Mhc-Drb* gene copy. Eighteen variable sites were detected among the 171 bp sequences, out of which eight corresponded to non-synonymous sites. Eight SNPs were genotyped over all other candidate genes: four SNPs appeared to be monomorphic out of the five SNPs analyzed within *Tlr4* and were excluded from the data. No polymorphism was found within *Tlr7.* Finally, polymorphism of a single SNP was analyzed for each of the following immune-related genes: *Mx2*, *Tnf* promoter and *Tlr4*.

### 3.2. Genetic diversity at local and global scales

A total of 43 pairs of microsatellites exhibited significant LD over the 684 tested (6%) after FDR correction. The loci involved differed across years and sites. Two out of 60 tests for deviation from HWE were significant, and both concerned the locus Cg16H5.

Supplementary Table S1 gives the range of diversity estimates, across years and sites, and at the whole metapopulation scale, for microsatellites markers and immune-related genes. Detailed results of the GLMs used to test the effects of bank vole population cycles on genetic diversity indices are presented in Supplementary Table S4. *H*_e_ and *N*_a_ estimates were higher at microsatellite loci than at immune-related genes at the local (within site - *H*_*e*_: *X*^*2*^_*2*_ = 45.69, *p* < 10^−4^; *N*_a_: *X*^*2*^_*2*_ = 25.72, *p* < 10^−4^, Fig. 2) and global scales (within metapopulation - *H*_e_ : *X*^*2*^_*2*_ = 169.20, *p* < 10^−4^; *N*_a_: *X*^*2*^_*2*_ = 19.48, *p* = 10^−4^, Fig. 2). At the local scale, *H*_e_ estimates also varied among years (*X*^*2*^_*2*_ = 8.47, *p* = 0.015), with lower estimates observed in 2008 than in 2007, and among sites (*X*^*2*^_*4*_ = 9.575, *p* = 0.048). At a global scale, *N*_a_ estimates varied among years (*X*^*2*^_*3*_ = 8.985, *p* = 0.029), with lower estimates observed in 2009 compared to 2005. We noted that the loss of alleles was of similar amplitude at microsatellites and *Mhc-Drb* (about two alleles lost out of 12 between 2005 and 2009, Table 1). Interestingly, one *Mhc-Drb* variant was lost during the crash between 2005 and 2007, and was not recovered at the metapopulation scale the years after. Contrastingly, new *Mhc*-*Drb* haplotypes were detected in particular sites of the metapopulation in 2008 (e.g. Mygl-DRB*15 in site M and Mygl-DRB*08 in site F). Immune SNPs exhibited strong temporal changes of diversity compared to other loci (*H*_e_; interaction *year * type of loci*: *X*^*2*^_*6*_ = 41.67, *p* < 10^−4^). In particular, *Mx2*_162 and *Tlr4*_1662 exhibited lower *H*_e_ levels in 2005 compared to 2009 (from 0.25 to 0.35 and 0.21 to 0.34 respectively), mainly as a result of an increase in frequency of the less common variant between these two years.

**Fig 2.**
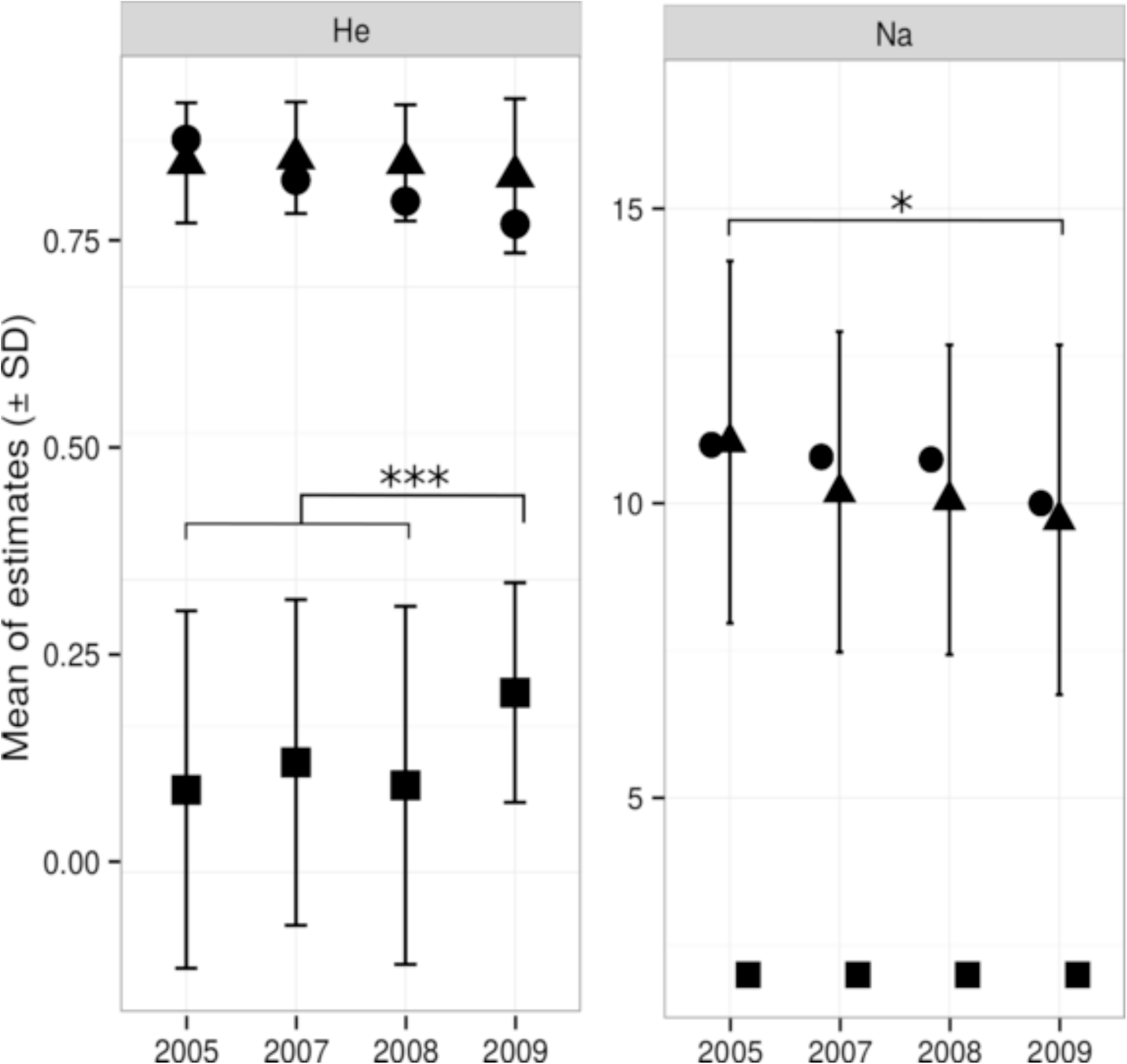
Temporal variations of *H*_e_ *and N*_a_ *e*stimates (mean ± SD) for the whole metapopulation, by type of locus. Mean *H*_e_ *and N*_a_ *e*stimates are represented as triangles for microsatellite loci, circles for *Mhc-Drb* and squares for SNPs. Bars represent standard deviations for microsatellite (15 loci) and SNPs (3 polymorphisms). A linear mixed model was performed with the four diversity indices estimated over the whole metapopulation sampled in 2005, 2007, 2008 and 2009. Significant differences between years are indicated by asterisks (* *p* < 0.05; *** *p* < 0.001).

### 3.3. Spatial genetic structure

Spatial pairwise *F*_ST_ estimates ranged between 0.003 and 0.056 for microsatellites, −0.077 and 0.270 for SNPs and −0.067 and 0.109 for *Mhc-Drb* (Table S5). At the scale of the metapopulation, temporal pairwise *F*_ST_ ranged between 0.003 and 0.017 at microsatellites, with high estimates corresponding to pairs of sampling date including the crash year 2009. None of these estimates were significant for immune-related genes (Table S6).

AMOVA analyses revealed significant spatial genetic structure at microsatellites only, and significant variation among sites within years for all loci, with higher percentage of variation detected at *Mhc-Drb* than at microsatellites or immune-related SNPs (Table 1A).

AMOVA analyses also revealed an absence of temporal genetic differentiation over the whole metapopulation whatever the loci considered, and significant variation among years within sites for *Mhc-Drb* and microsatellites, with higher percentage of variation detected at *Mhc-Drb*. No such local temporal genetic changes were detected at immune-related SNPs (Table 1B).

**Table 1.**
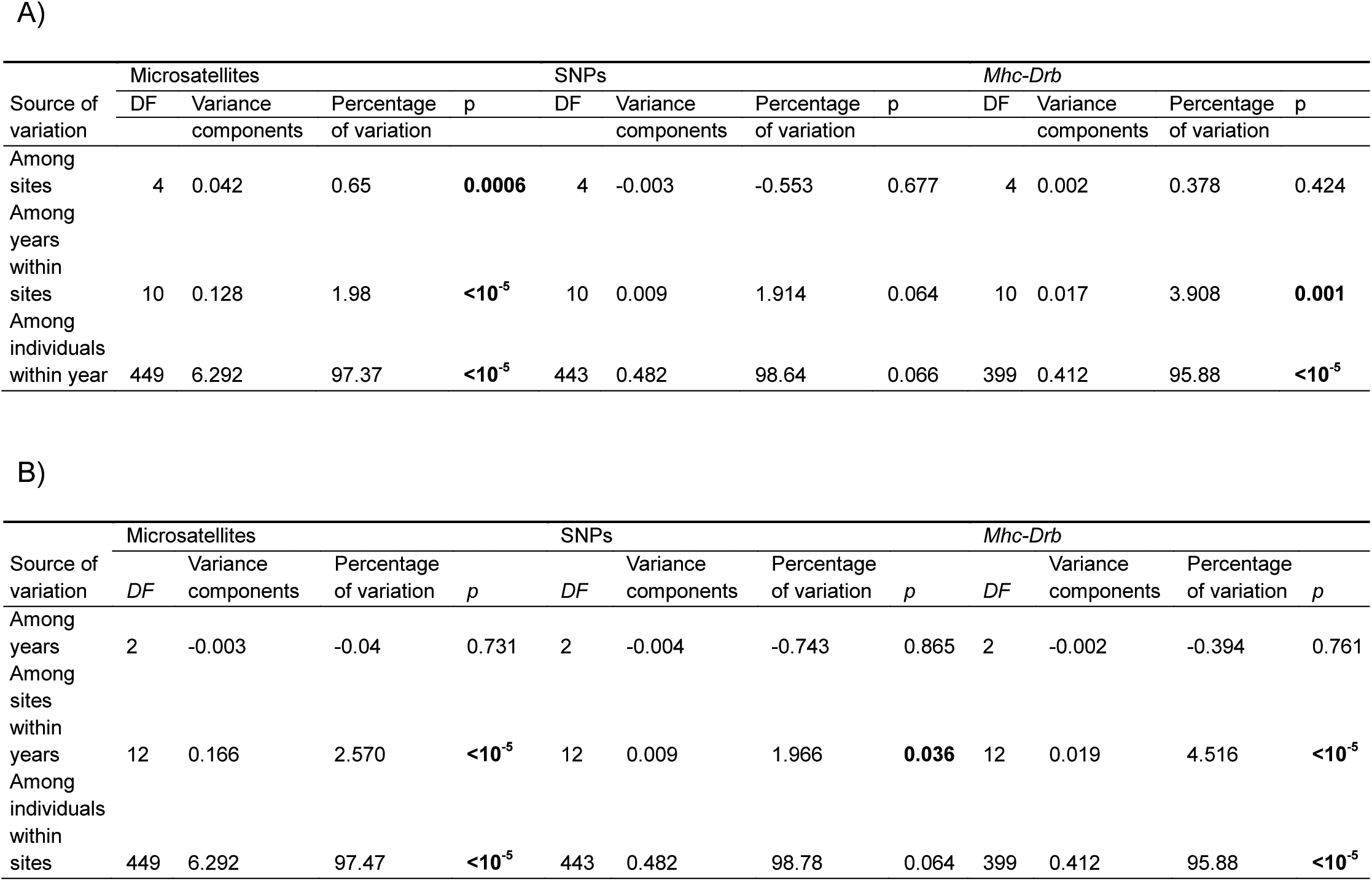
Analysis of molecular variance (AMOVA) were performed considering the five sites B, C, F, G, O sampled in 2005, 2007 and 2008 (sample size larger than 8 individuals) and the three types of loci, *i.e.* microsatellites, immune SNPs and *Mhc-Drb*. Two hierarchies were examined: A) sites as groups and years as subdivisions within sites; B) years as groups and sites as subdivisions within years.

Clustering analyses based on microsatellites performed over the whole dataset revealed three clusters that did not discriminate either years or sites. All voles were assigned to these three genotypic groups (Fig 3A, B). It is therefore likely that a complete absence of spatiotemporal structure (*K* = 1) is the best hypothesis to explain this result.

**Fig 3.**
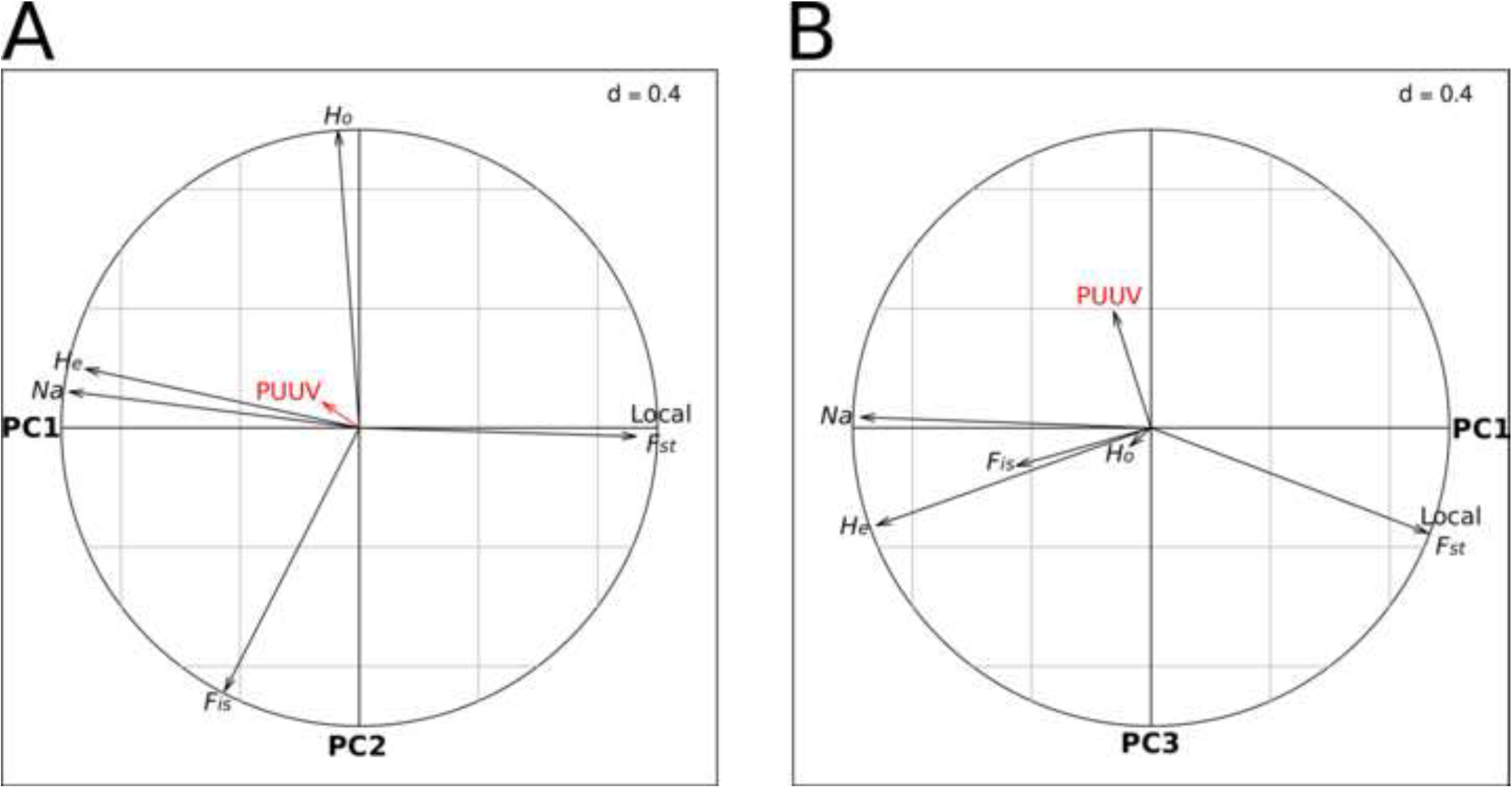
STRUCTURE plots of the bank vole metapopulation over the whole survey. The figure presents A) the bar plots resulting from the clustering analysis based on microsatellites and performed for *K* = 3 on bank voles sampled over the whole survey. Within bar plots, each individual is represented by a single vertical line broken into *K* colored segments, which lengths are proportional to the individual membership coefficients, in each inferred cluster. Each color represents a different genotypic group. B) The rate of change in the log probability of the data between successive *K* values (Δ(*K*) *i.e.* L”(*K*) divided by the standard deviation of L(*K*)) is shown for *K* = 2 to *K* = 9.

Spatial autocorrelograms revealed a strong decrease of relatedness between individuals with geographic distance, whatever the sex and year considered (Fig 4). Nevertheless, mean relatedness of females was slightly higher at very small spatial scale (within sampling site or in the first surrounding sites distant from less than 1 km) during peak years (respectively 0.04 and 0.05 in 2005 and 2008) than during the increase phase (0.03 in 2007). Contrastingly, relatedness between males was higher within sites during the increase phase (0.03) than during peak years (respectively 0.01 and 0.02 in 2005 and 2008).

**Fig 4.**
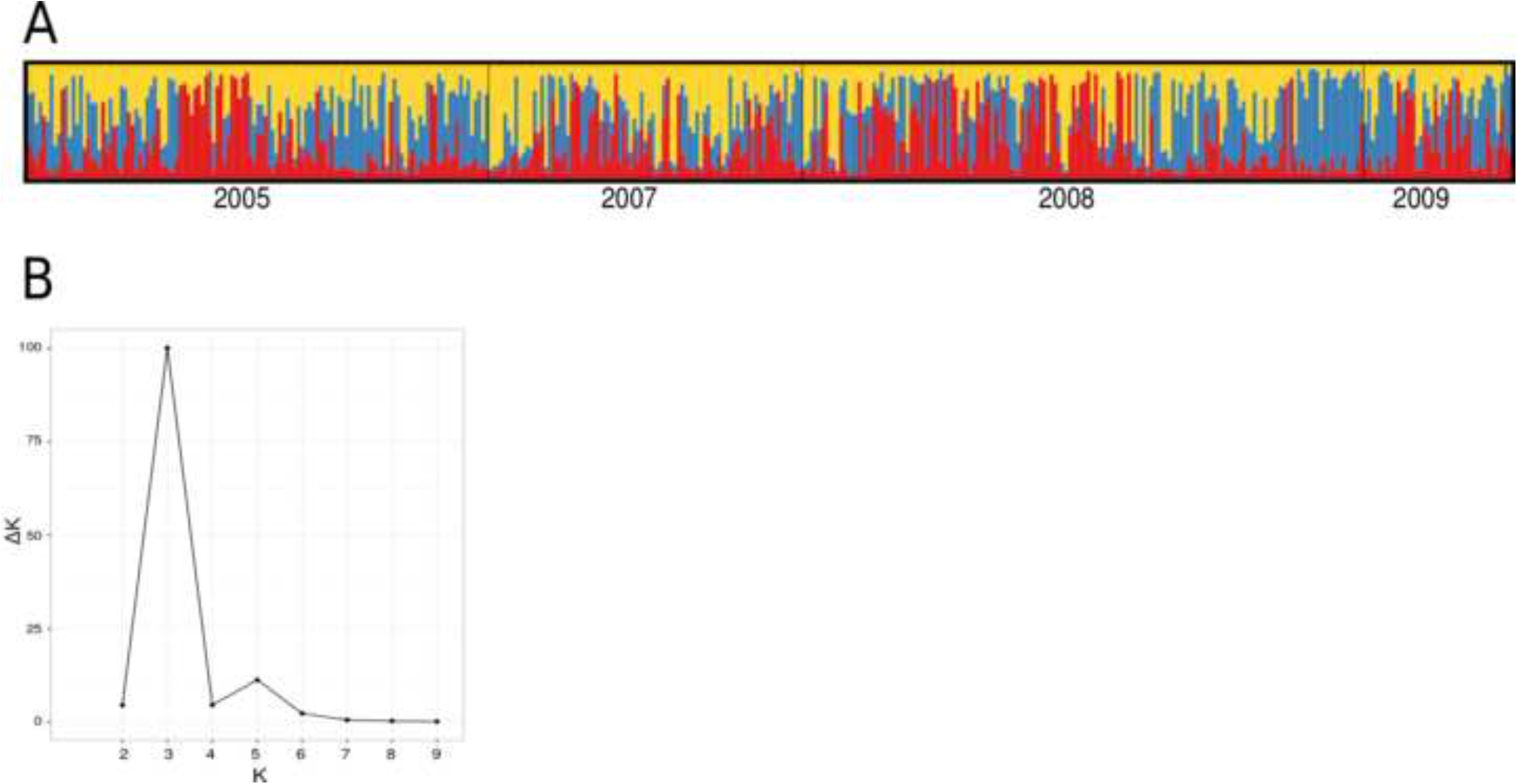
Spatial autocorrelation analyses based on Li’s relatedness coefficient. Correlograms are presented independently for males and females and for each year of sampling between 2005 and 2008. Geographic distances classes are indicated in km. The mean and standard error of Li’s relatedness coefficient (Li et al., 1993) are indicated for each class.

### 3.4. Links between genetic structure and PUUV epidemiology

For each year, the estimates of mean relatedness among seropositive and seronegative voles in the metapopulation showed very low and similar values, respectively *r* = −0.006 and *r* = 6.10^−4^ in 2005 (*p* = 0.344), *r* = −0.014 and *r* = −0.006 in 2007 (*p* = 0.857), *r* = −0.009 and *r* = 0.005 in 2008 (*p* = 0.333) and *r* = −0.050 and *r* = 0.030 in 2009 (*p* = 0.399).

Site-specific *F*_ST_ estimates at microsatellite markers (computed using the GESTE software) ranged between 0.009 and 0.063, showing various levels of genetic isolation of sites within the bank vole metapopulation (Table S1). The first axis of the PCA based on microsatellite genetic parameters (PC1) explained 57.4% of the variability and revealed a strong opposition between these local *F*_ST_ estimates and the other genetic features describing genetic diversity, especially *H*_e_ and *N*_a_ (Fig 5A). This axis therefore illustrated the opposing impacts of genetic drift and migration on local genetic diversity. The second axis of the PCA (PC2) accounted for 36.5% of the total variability, and opposed microsatellite genetic parameters reflecting the deviation from Hardy-Weinberg equilibrium within site (*H*_o_ and *F*_is_). The third PCA axis (PC3) explained 5.1% of the variability and contrasted the diversity indices *N*_a_ and *H*_e_ (Fig 5B). The projection of the variable *year* on this axis revealed that PC3 discriminated the increase phase 2007 after the decline in 2006 and peak years (2005, 2008). The best and the most parsimonious model explaining PUUV seroprevalence using site coordinates on these three axes included PC3 only (*z*-value = 2.119, *p* = 0.034). PUUV prevalence therefore seemed mostly related to local changes in microsatellite allelic richness associated with the population cycle, PUUV seroprevalence reaching higher levels during the increase year (2007).

**Fig 5.**
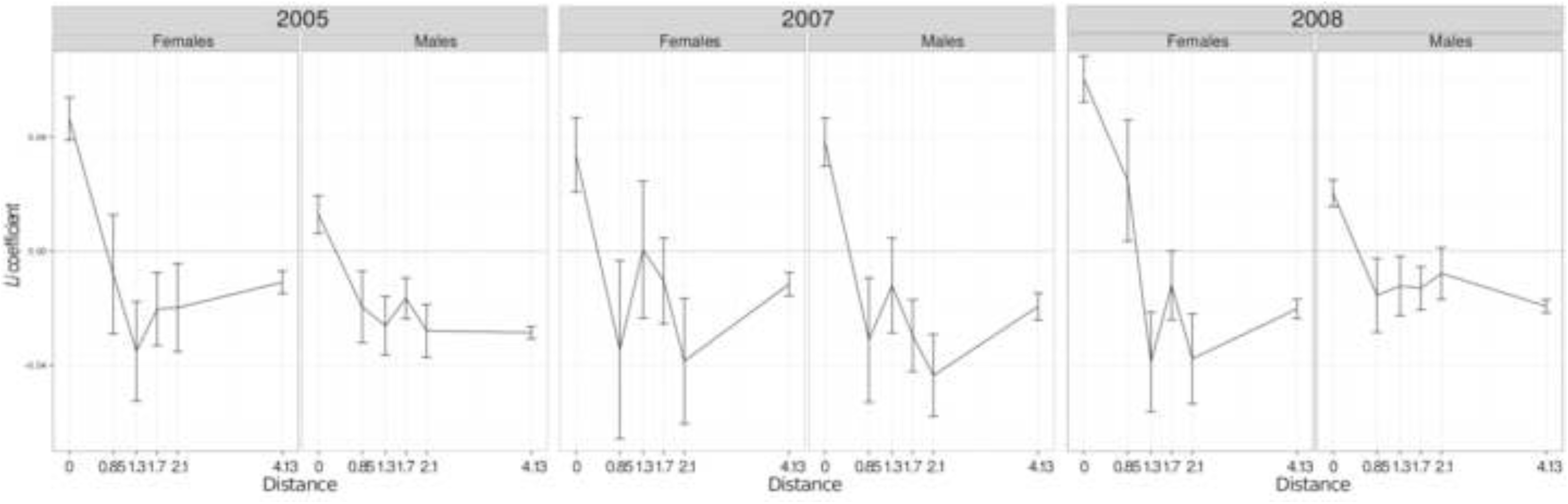
Principal component analysis based on local genetic parameters estimated using microsatellites. Correlation circles of the five genetic variables using A) PC1 and PC2, and B) PC1 and PC3. Puumala seroprevalence was included as a supplementary variable.

### 3.5. Detection of signatures of selection

The patterns of differentiation conditional on heterozygosity differed between genes and years (Fig 6). Departure from neutral expectation was observed at the *Tlr4*-1662 SNP in 2008 only. *F*_ST_ estimate lay outside the 95% confidence region of the null distribution, showing that the *Tlr4*-1662 SNP was more differentiated (*F*_ST_ = 0.076) than expected under neutrality (Fig 6C), showing a significant signature of positive selection (*p* = 0.037). *Mx2*-162 SNP exhibited low *F*_ST_ estimates (*F*_ST_ = −0.030 in 2007; *F*_ST_ = −0.013 in 2008), suggesting potential signature of balancing selection (*p* = 0.090). No significant departure from neutral expectations was found for *Tnf* and *Mhc-Drb*. Investigating to what extent the approach to detect signatures of selection was robust to temporal variations in population size, we found virtually no difference between the distributions of *F*_ST_ conditional on heterozygosity across different scenarios of population size changes (see Supplementary Material Fig. S3), which justifies *a posteriori* the use of our approach to detect signatures of selection on cyclic populations.

**Fig 6.**
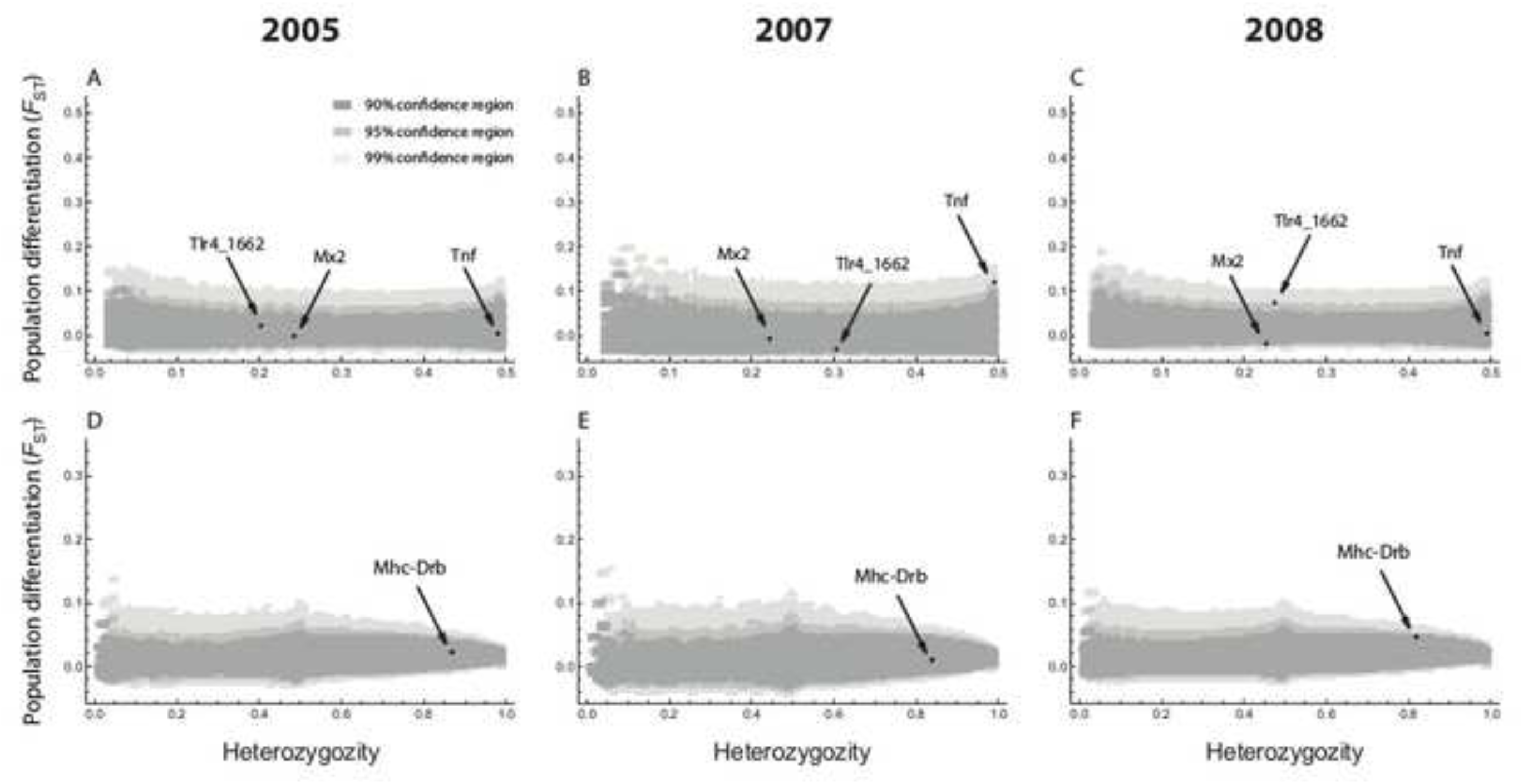
Distribution of *F*_ST_ conditional on the overall heterozygosity (*H*_e_) at immune-related genes, for the whole metapopulation dataset in 2005, 2007 and 2008. Black dots represent the observed, locus-specific, estimates of differentiation and heterozygosity for SNPs in candidate genes (A-C) and for *Mhc-Drb* (D-F). The 90%, 95% and 99% confidence regions of the null distribution (obtained by means of stochastic simulations in a neutral model) are shown in dark grey, medium grey and light grey, respectively. The density was obtained using the average shifted histogram (ASH) algorithm (Scott, 2002) with smoothing parameter *m* = 2.

### 3.6. Associations between immune-related genotypes and PUUV

Immune gene polymorphism did not globally discriminate PUUV seropositive and seronegative voles (*p* = 0.077). Five alleles of the *Mhc-Drb* gene tended to be associated with PUUV serological status. The RR values of Mygl-DRB*106, Mygl-DRB*08 and Mygl-DRB*94 reached respectively 1.724, 1.717 and 1.464, indicating that individuals carrying this allele were almost twice more likely to be PUUV seropositive than others (Fig 7A,B). On the contrary, with a respective RR value of 0.263 and 0.339, bank voles carrying Mygl-DRB*105 and Mygl-DRB*15 were more than twice less likely to be PUUV seropositive than others (Fig 7A,B). After Bonferroni sequential corrections, these RR values were not significant. Note that similar results were observed when considering only over-wintered bank voles that had a long time of potential exposure to PUUV (results not shown).

**Fig 7.**
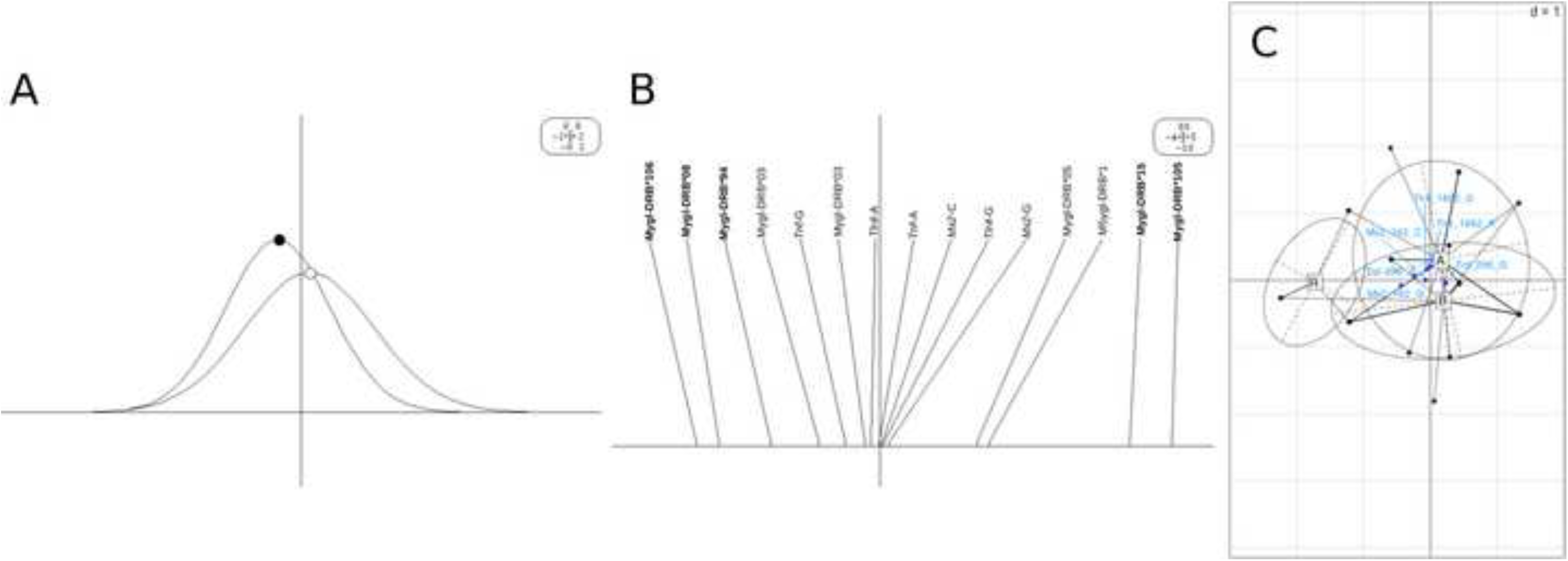
Discriminant analysis performed on immune-related polymorphism of PUUV adult bank voles over the whole metapopulation. A) PUUV seropositive adult bank voles, represented by black dot, are marginally different from PUUV-seronegative ones, represented by white dot. B) Alleles indicated in bold (Mygl-DRB*106, Mygl-DRB*08, Mygl-DRB*94, Mygl-DRB*15 and Mygl-DRB*105) are associated with PUUV serological status. C) Representation of the two first axes of the within/between analysis performed on seropositive adult bank voles only, with A, B, R corresponding to the PUUV genogroups defined by Razzauti et al. (Razzauti et al., 2013, 2008) and immune-related gene polymorphisms indicated by blue dots.

We found a marginally significant association between bank vole polymorphism at *Mx2*-162 SNP (presence/absence of G) and PUUV genogroup. Individuals carrying the G nucleotide at *Mx2*-162 SNP were at higher risk of being infected with PUUV reassortants (S segment, genogroup A & M segment genogroup B or S segment, genogroup B & M segment genogroup A; RR = 7.500, *p* = 0.052) than those carrying the C base only. The within/between analysis also suggested that this *Mx2*-162 SNP could be important with regard to PUUV (Fig 7C).

## 4. Discussion

In this study, we analysed the genetic variation of a bank vole metapopulation, experiencing three-year population density cycles, at microsatellite loci and immune-related genes to examine the potential interactions between microevolutionary processes and PUUV epidemiology. Overall, we showed that neutral and adaptive genetic variability was maintained through years despite repeated demographic bottlenecks. Rapid increase in vole density, the absence of landscape barriers and high gene flow seemed to counterbalance the effects of genetic drift, which may explain the absence of founder effects observed in PUUV population throughout vole population cycles. In addition, our results suggest that PUUV circulation results from vole dispersal through sex-specific and density-dependent movements, but not from contacts between relatives. Finally, we found little evidence of selection acting on immune-related genes within this metapopulation. We characterized departure from neutral expectations at *Tlr4* gene in 2008 only, and marginal associations between PUUV genetic variants and bank vole alleles at *Mhc-Drb* and *Mx2*). Microevolutionary changes and PUUV epidemiology in this metapopulation therefore seemed to be mainly driven by neutral processes.

### 4.1. Genetic drift, bank vole dispersal and PUUV epidemiology

In boreal Fennoscandia, bank vole populations experience strong multiannual fluctuations in abundance, with peaks occurring every three to five years and abundance varying by a factor 500 between low and peak phases (Hansson and Henttonen, 1988). Such regular and severe declines in population size are expected to affect neutral genetic variation through genetic drift. However, several studies have shown that cyclic rodent populations maintained high levels of genetic diversity, at least at a metapopulation scale (Berthier et al., 2005; Ehrich et al., 2009, 2001; García-Navas et al., 2015; Gauffre et al., 2014; Winternitz et al., 2014). Marginal and temporal decreases in allelic richness were previously observed by Rikalainen et al. (2012), who studied Finnish bank vole populations at a larger scale (100 km^2^) around Konnevesi. Our results provided similar patterns, with high levels of heterozygosity and allelic richness being observed whatever the cycle phase considered. We detected a decrease in allelic richness in the crash phase of 2009 at the scale of the metapopulation. This loss of diversity might be only transient as no decrease could be observed after the 2006 crash year. Therefore, fluctuations in population size did not lead to strong signature of bottleneck in the genetic data, whatever the spatial scale considered. Similarly, Razzauti et al. (2013) did not find any evidence of founder effects affecting PUUV diversity in the area including this vole metapopulation. Significant loss of PUUV variants were detected in 2009 only, with the complete absence of variants from one of the two PUUV genogroups otherwise detected since 2005. These patterns could be explained by the low sampling size in 2009. But it is likely that the low phase is usually too short to impact host genetic diversity, and/or that strong bank vole gene flow might compensate for genetic drift (Ehrich et al., 2009). Genetic patterns of temporal and spatial differentiation supported the hypothesis that dispersal among sites may rapidly counterbalance the potential loss of genetic diversity experienced by vole and PUUV populations. Although significant temporal genetic differentiation was locally detected between years, low estimates of temporal genetic differentiation were detected at the scale of the metapopulation. These estimates were slightly higher when comparing pairs of sampling dates before and after decline (especially in 2009). This result corroborated the transient impact of genetic drift on the distribution of genetic variability, and the importance of gene flow occurring after population crash, once vole population size increased again. The relative temporal genetic stability of this metapopulation was also supported by the clustering analysis performed over the whole survey, as it did not reveal any consistent spatial structuring. Altogether, these results emphasized the absence of environmental heterogeneity within the studied area, a typical boreal forest, that could disrupt bank vole gene flow and in turn, PUUV dispersal among sites. The possibility for voles to move all around this area provides an explanation for the absence of spatial structure detected in PUUV population across this area (Razzauti et al., 2013).

Spatial autocorrelation analyses highlighted changes in the bank vole spatial genetic structure from year to year, that probably resulted from density fluctuations favouring dispersal and population turnover (e.g. Ehrich et al., 2009; Rikalainen et al., 2012)). This raised the question of density-dependent dispersal, that is, whether dispersal rate increased during low phase or with increasing population size (see Pilot et al. (2010) for references and predictions). As evidenced by Rikalainen et al. (2012) and Gauffre et al. (2014), our study suggested patterns of sex-specific and density-dependence dispersal. Negative density dependent migration of females could underlie the local changes in relatedness observed between low density and peak years. The slightly lower relatedness observed during the increase phase could hence be explained by the colonisation of new empty sites by females. The high levels of relatedness observed during 2005 and 2008 could reflect female phylopatry and low migration (see Le Galliard et al. 2012) once population size has reached high abundance. On the other hand, the lower levels of relatedness observed in peak years compared to the increase phase when focusing on males could result from their high dispersal rate and high gene flow once density reached high levels. The social behavior in *M. glareolus* is likely to underlie these variations of dispersal through time. Dispersal is strongly connected with maturation, which is season and density-dependent. Maturation phases and connected dispersal of young maturing voles are shorter in the peak than in the increase summer, as they only concern the first litter produced in spring. This sex-biased density dependent migration could be of main importance in shaping PUUV epidemiology. Indeed, a male bias in PUUV infection was detected in Konnevesi in overwintered, breeding bank voles, whereas a female bias was seen in summer-born breeding animals probably as a result of more frequent contact with old (infected) males or aggressive encounters with other breeding females during the territory establishment (Voutilainen et al., 2016). In addition, PUUV seropositive voles were found to be more abundant during the high-density years of the dynamic cycle (increase and peak autumn - winters), as confirmed by the strong correlation observed in this study between PUUV seroprevalence and two indices of genetic diversity differentiating peak *versus* low-density years. We could therefore suggest that during low-density phase, female voles could play a role in the colonization of empty sites and consequently in the transmission of PUUV. During peak years, males would be responsible for repeated dispersal events and gene flow, therefore contributing to PUUV transmission.

### 4.2. Impact of metapopulation dynamics on immune-related gene polymorphism

Whether migration and natural selection can counterbalance the effect of drift on immune gene polymorphism in populations experiencing drastic fluctuations remains an open research topic, as empirical data have revealed contrasted patterns. From an eco-evolutionary perspective, this question is of uttermost importance as a loss of genetic variation could result in dampened adaptive capability, which is especially problematic when faced with rapidly evolving pathogens. A recent meta-analysis showed that a greater loss of genetic diversity was generally seen at *Mhc* genes rather than at neutral loci, potentially because of the uneven distribution of haplotypes before the bottleneck (Sutton et al., 2011). Here, we studied five genes contributing to immunity, with polymorphism analyzed within protein coding regions (*Mhc-Drb*, *Tlr*s, *Mx2*) or in the promoter (*Tnf*-□).Variation at these immune-related genes has previously been shown to be associated with rodent responses to PUUV infections (Charbonnel et al., 2014).

Our study first enabled to investigate the relative role of neutral processes and selection in shaping the geographic variation of immune gene diversity in this cyclic bank vole metapopulation. As predicted under genetic drift only, reduction of immune-related gene variability was observed at *Mhc-Drb* gene during the crash year 2009, and it was of similar amplitude than the decrease observed at microsatellites. We also noted a particular allele present at low frequencies in one site during the 2005 peak year could be lost and never recovered within the metapopulation after the 2006 crash. Such impact of genetic drift was also detected on SNPs, with different patterns detected according to the immune-related gene considered. Loss of less frequent variants was for example locally observed after the 2006 crash at *Mx2*_162 SNP only. However, this impact of genetic drift on immune-related genes seemed minor and very transient, as there were no such significant temporal changes in allelic frequencies at *Mhc*-*Drb* or other SNPs. This contrasts with the conclusion drawn from the meta-analysis of Sutton et al. (2011). They proposed that the greater reduction in *Mhc* gene diversity compared to neutral genetic diversity may be observed when negative frequency-dependent selection acting before bottlenecks results in the uneven distribution of most *Mhc* variants/haplotype, accelerating the loss of *Mhc* variability.

Dispersal can be invoked to explain that allelic diversity was recovered once vole abundance increased, as previously seen in Winternitz et al. (2014). The detection of previously unseen *Mhc*-*Drb* haplotypes in particular sites of the metapopulation during the 2008 peak year corroborated this possibility, assuming that it resulted from the immigration of bank voles from outside the metapopulation, carrying these different haplotypes.

### 4.3. Little evidence of selection acting on immune-related gene polymorphism

Model-based simulations revealed little evidence of departures from neutrality for immune-related gene polymorphisms. The *Tlr4*-1662 SNP showed signature of positive selection, in 2008 only. *Tlr4* is part of *Tlrs* gene family and is involved in coevolutionary processes with a wide variety of pathogens (Akira et al., 2006; Hughes and Piontkivska, 2008). Several studies recently emphasized the impact of historical and contemporary positive selection on *Tlr4* gene polymorphism in rodents (Fornůsková et al., 2014, 2013; Turner et al., 2012), and it is likely that pathogens might mediate this selection. The signature of positive selection observed in this study might reflect spatial heterogeneity in the whole pathogen community or in one/few pathogens recognized by TLR4. Temporal variations of selection patterns at immune-related genes have previously been described in arvicoline rodents experiencing population cycles. Bryja et al. (2007) found density-dependent changes in selection pressure throughout a complete density cycle of the montane water vole (*Arvicola scherman*), and these changes were detected at only one *Mhc* class II gene over the two examined. In cyclic populations of field vole (*Microtus agrestis*), Turner et al. (2012) detected signature of selection acting on only two genes encoding cytokines over the twelve immune-related genes examined. Such results could be mediated by temporal heterogeneity in pathogen pressure (e.g. Fraser et al., 2010) that may have a strong selective effect even in a few generations (e.g. Oliver and Piertney, 2012). Because positive selection was observed in the peak year, when gene flow is important once voles get mature in spring and should homogenize directly transmitted pathogens, potentially other candidates than PUUV should preferentially be searched among vector-borne agents (e.g. bacteria like *Bartonella* sp., *Borrelia* sp.) or pathogens able to survive outside their hosts (e.g. ectoparasites).

A marginally significant pattern of balancing selection was observed at *Mx2* gene in 2008 and we found a tendency for PUUV reassortants to be associated with a particular *Mx2* genotype. *Mx2* gene encodes for proteins that provide immunity against viruses replicating in the cytoplasm, including hantaviruses (Jin et al., 2001; Lee and Vidal, 2002). The SNP detected under selection and associated with PUUV variants was located within the leucine zipper motif of the C-terminal GTP effector domain, where mutations can affect the antiviral efficacy of Mx proteins (Janzen et al., 2000; Kochs and Haller, 1999; Zürcher et al., 1992). Balancing selection could be mediated by spatiotemporal heterogeneity in candidate viruses including PUUV (Guivier et al., 2014; Razzauti et al., 2013, 2008), and it would be stronger during peak year when dispersal may homogenize their distribution within the area.

Finally, we did not confirm the signatures of positive selection that were previously detected at the −296 SNP of *Tnf*- promoter between NE endemic and non-endemic areas, at regional and European geographical scales (Guivier et al., 2014, 2010). Several explanations can be evoked to explain the absence of footprint of selection observed in this study. In particular, the statistical methods and the sampling design used in this study might prevent from detecting local adaptation mediated by pathogens (Lotterhos and Whitlock, 2014). First, some of the underlying assumptions of our tests to detect departures from neutral expectation are likely to be violated in this study. In particular, FDIST2 and the modified DFDIST are based on an island model at migration-drift equilibrium, while the true demographic history of *M. glareolus* is characterized by non-equilibrium population dynamics. However, we have shown by means of simulations (see Supplementary Appendix S1 and Supplementary Fig. S3) that the approach was robust to temporal variations in population size, at least in the range of differentiation level observed in our data, which is consistent with the idea that the distribution of *F*_ST_ estimates should be robust to the vagaries of demographic history (Beaumont, 2005). Second, *F*_ST_ outlier tests have low statistical power to detect balancing selection (Beaumont and Balding, 2004) and weak divergent selection (e.g. (Wachowiak et al., 2009). The spatial scale examined in this study might be too small to cover different selective environments mediated by pathogens and to lead to strong effective selection. It would therefore be interesting to perform similar analyses at a larger scale (about 100 km^2^) to cover contrasted ‘pathogenic landscapes’ but still limiting the potential confounding effects due to neutral population structure. Finally, we can not discard the possibility that we may have missed important immune related genes evolving under strong effective selection in this metapopulation.

In conclusion, this study provided evidence for the role of vole dispersal on PUUV circulation through sex-specific and density-dependent movements. We also revealed that multiannual dynamic cycles were likely to affect the diversity of immune-related genes involved in susceptibility to PUUV, mainly through neutral microevolutionary processes. In the future, we recommend both the development of population genome-wide scan approaches that require no *a priori* on immune-related candidate genes and the analysis of metapopulations covering heterogeneous landscapes with regard to pathogens, to better assess the influence of selection on rodent/hantavirus interactions. Preliminary high-throughput characterization of rodent microbial communities (e.g. Galan et al., 2016) should enable to define the optimal geographical scale to consider for such studies, *i.e.* large enough to encompass contrasted environments with regard to pathogen community diversity and structure, but still limited to remain close from an island model of demographic history.

## Acknowledgments

Microsatellite data used in this work were produced through the technical facilities of the Centre Méditerranéen Environnement Biodiversité (CeMEB). We are grateful to Audrey Rohfritsch and Viola Walther for the identification of SNPs at immune gene during the pilot study. We are grateful to Jeremy Gaudin for technical assistance and preliminary analyses.

This research has been partially funded by EU grants GOCE-CT-2003-010284 EDEN and FP7-261504 EDENext, and the paper is catalogued by the EDENext Steering Committee as EDENext 405 (http://www.edenext.eu). AD is currently funded by an INRA-EFPA / ANSES fellowship.

## Supporting Information

**S1 Protocol:** SNPs identification and genotyping

**S1 Table**: Characteristics and genetic diversity estimates within each year and each sampling site, including the number of bank voles sampled (N) and PUUV positives (Ni), PUUV seroprevalence and the genetic diversity estimates (*H*_*o*_, *H*_*e*_, *F*_*IS*_, *N*_a_). When less than eight individuals were sampled in a site, local estimates were not calculated (-and all the sites in 2009). ‘Total’ refers to the whole metapopulation. Significant values are in bold.

**S2 Table**: Analyses of single nucleotide polymorphism for four immune-related genes. A) Primers used for *M. glareolus* identification, amplification of gDNA for PCR and sequencing with corresponding fragment length and total length of the sequence. B) Polymorphism for three immune-related genes screened using a sample of 12 to 29 individuals from three European bank vole populations. Singletons (polymorphism was detected in only one heterozygote indivduals) were not included in further genotyping analyses of Konnevesi populations. The software PolyPhen (http://genetics.bwh.harvard.edu/pph; Ramensky et al. 2002) was used to predict the functional impact of non-synonymous SNPs on the translated protein, based on biochemical and physical characteristics of the amino acid replacements.

**S3 Table**: Cycling conditions of PCRs.

**S4 Table**: Results of the two linear mixed models. Only significant pairwise interactions are reported.

**S5 Table**: *F*_ST_ estimates between sites per year (lower side) and associated *p*-values of Fisher exact tests (upper side).

**S6 Table**: *F*_ST_ estimates between years over the whole metapopulation (lower side) and associated p-values of Fisher exact tests (upper side).

**S7 Table**: This file includes A) the individual and spatial data of all bank voles, B) the microsatellite genotypes for 19 loci, C) the genotypic data for four immune-related genes (eight independent SNPs genotyped using the KASPar technology) and D) the *Mhc-Drb* exon 2 genotypes (sequences are available in Genbank).

**S1 Figure**: Distribution of *F*_ST_ conditional on the overall heterozygosity (*H*_e_) at microsatellites, for the whole metapopulation dataset in 2005, 2007 and 2008. Black dots represent the observed, locus-specific, estimates of differentiation and heterozygosity for microsatellites. The 90%, 95% and 99% confidence regions of the null distribution (obtained by means of stochastic simulations in a neutral model) are shown in dark grey, medium grey and light grey, respectively. The density was obtained using the Average Shifted Histogram (ASH) algorithm (Scott, 2002) with smoothing parameter *m* = 2.

**S2 Figure**: Phylogenetic relationships of bank vole 454 (202 bp)-DRB sequences (this study; Mygl-DRB*120–151) and previously known Cluster I, Cluster II and Group III bank vole sequences from (Axtner and Sommer, 2007); Clgl-DRB*01-28 (eliding the numbers 2 and 9); EF434791– EF434816] expressed as neighbour-joining tree based on Kimura’s two-parameter distances, pairwise deletion. Bootstrap values are based on 1,000 replicates. Species specific Mhc nomenclature is used. H2 IEb (Mus musculus DRB: U28489.1) serves as outgroup.

**S3 Figure**: Distribution of *F*_ST_ conditional on the overall heterozygosity (*H*_e_) at immune-related genes, for the whole metapopulation dataset in 2007 and 2008, for different demographic scenarios: constant deme size (A-B), cyclic deme size (C-D) and randomly-varying deme size (E-F). See Supplementary Appendix S1 for further details on the simulations. Black dots represent the observed, locus-specific, estimates of differentiation and heterozygosity for SNPs in candidate genes. The 90%, 95% and 99% confidence regions of the null distribution (obtained by means of stochastic simulations in a neutral model) are shown in dark grey, medium grey and light grey, respectively. The density was obtained using the average shifted histogram (ASH) algorithm (Scott, 2002) with smoothing parameter *m* = 2.

**S1 Appendix:** This file includes details on the simulations performed to investigate to what extent the modified DFDIST approach to detect signatures of selection was robust to temporal variations in population size.

